# Tissue-specific and interferon-inducible expression of non-functional ACE2 through endogenous retrovirus co-option

**DOI:** 10.1101/2020.07.24.219139

**Authors:** Kevin Ng, Jan Attig, William Bolland, George R. Young, Jack Major, Andreas Wack, George Kassiotis

## Abstract

Angiotensin-converting enzyme 2 (ACE2) is an entry receptor for Severe Acute Respiratory Syndrome Coronavirus 2 (SARS-CoV-2), as well as a regulator of several physiological processes. *ACE2* has recently been proposed to be interferon-inducible, suggesting that SARS-CoV-2 may exploit this phenomenon to enhance viral spread and questioning the efficacy of interferon treatment in Coronavirus disease 2019 (COVID-19). Using a recent *de novo* transcript assembly that captured previously unannotated transcripts, we describe a novel isoform of *ACE2*, generated by co-option of an intronic long terminal repeat (LTR) retroelement promoter. The novel transcript, termed *LTR16A1-ACE2*, exhibits specific expression patterns across the aerodigestive and gastrointestinal tracts and, importantly, is highly responsive to interferon stimulation. In stark contrast, expression of canonical *ACE2* is completely unresponsive to interferon stimulation. Moreover, the *LTR16A1-ACE2* translation product is a truncated, unstable ACE2 form, lacking domains required for SARS-CoV-2 binding and therefore unlikely to contribute to or enhance viral infection.

## Introduction

Interferons represent the first line of defence against viruses in humans and other jawed vertebrates (Sadler and Williams, 2008). Recognition of viral products in an infected cell results in autocrine and paracrine signalling to induce an antiviral state characterized by expression of a module of interferon-stimulated genes (ISGs) that restrict viral replication and spread (Sadler and Williams, 2008; Stetson and Medzhitov, 2006). Indeed, recombinant interferon is often given as first-line therapy in viral infection (Gibbert et al., 2013) and preliminary results suggest that interferon treatment may be effective against Coronavirus disease 2019 (COVID-19) (Hung et al., 2020; Wang et al., 2020).

Interferon signalling results in rapid upregulation of several hundred ISGs, including genes that inhibit various stages of viral entry and replication as well as transcription factors that further potentiate the interferon response (Sadler and Williams, 2008; Stetson and Medzhitov, 2006). Given that unchecked interferon signalling and inflammation can result in immunopathology, ISGs are subject to complex regulatory mechanisms (Ivashkiv and Donlin, 2014).

At the transcriptional level, long terminal repeats (LTRs) derived from endogenous retroviruses and other LTR-retroelements serve as cis-regulatory enhancers for a number of ISGs and are required for their induction (Chuong et al., 2016). Adding to this regulatory complexity, many LTRs are themselves promoters responsive to interferon and are upregulated following viral infection or in interferon-driven autoimmunity (Attig et al., 2017; Tokuyama et al., 2018; Young et al., 2012; Young et al., 2014).

The co-evolution of viruses and hosts has resulted in a number of strategies by which viruses evade or subvert interferon responses (García-Sastre, 2017). Compared with other respiratory viruses, Severe Acute Respiratory Syndrome Coronavirus 2 (SARS-CoV-2) elicits a weak interferon response despite strong induction of other chemokines (Blanco-Melo et al., 2020). Though the mechanism by which SARS-CoV-2 dampens interferon responses remains unclear, the ORF3b, ORF6, and nucleoprotein of the closely-related SARS-CoV function as interferon antagonists (Kopecky-Bromberg et al., 2007). SARS-CoV-2 uses angiotensin-converting enzyme 2 (ACE2) as its primary receptor (Hoffmann et al., 2020; Shang et al., 2020) and recent work suggested that SARS-CoV-2 may hijack the interferon response by inducing *ACE2* expression (Ziegler et al., 2020). By integrating multiple human, macaque, and mouse single-cell RNA-sequencing (RNA-seq) datasets, Ziegler et al. identified *ACE2* as a primate-specific ISG upregulated following viral infection or interferon treatment (Ziegler et al., 2020). Use of an ISG as a viral receptor would result in a self-amplifying loop to increase local viral spread, and calls into question the efficacy and safety of recombinant interferon treatment in COVID-19 patients.

Using our recent *de novo* transcriptome assembly (Attig et al., 2019), we identify a novel, truncated *ACE2* transcript, termed *LTR16A1-ACE2*, initiated at an intronic *LTR16A1* retroelement that serves as a cryptic promoter. Strikingly, we find that the truncated *LTR16A1-ACE2* and not full-length *ACE2* is the interferon-inducible isoform, and is strongly upregulated in viral infection and following interferon treatment. Importantly, the protein product of the *LTR16A1-*ACE2 transcript does not contain the amino acid residues required for SARS-CoV-2 attachment and entry and is additionally post-translationally unstable. These findings have important implications for the understanding of *ACE2* expression and regulation, and thus for SARS-CoV-2 tropism and treatment.

## RESULTS

### *LTR16A1-ACE2* is a tissue-specific novel isoform of *ACE2*

Our recent *de novo* cancer transcriptome assembly (Attig et al., 2019) identified a chimeric transcript between annotated exons of *ACE2* and an *LTR16A1* retroelement, integrated in intron 9 of the *ACE2* locus. This transcript, which we refer to here as *LTR16A1-ACE2*, is initiated at the *LTR16A1* retroelement, which functions as a cryptic promoter and includes exons 10-19 of *ACE2* (Figure 1A). Promoter activity of the *LTR16A1* retroelement was highly supported by splice junction analysis of RNA-seq data from TCGA lung adenocarcinoma (LUAD) and lung squamous cell carcinoma (LUSC) (Figure 1A) and by promoter-based expression analyses of the FANTOM5 data set (Figure S1).

**Figure 1.**
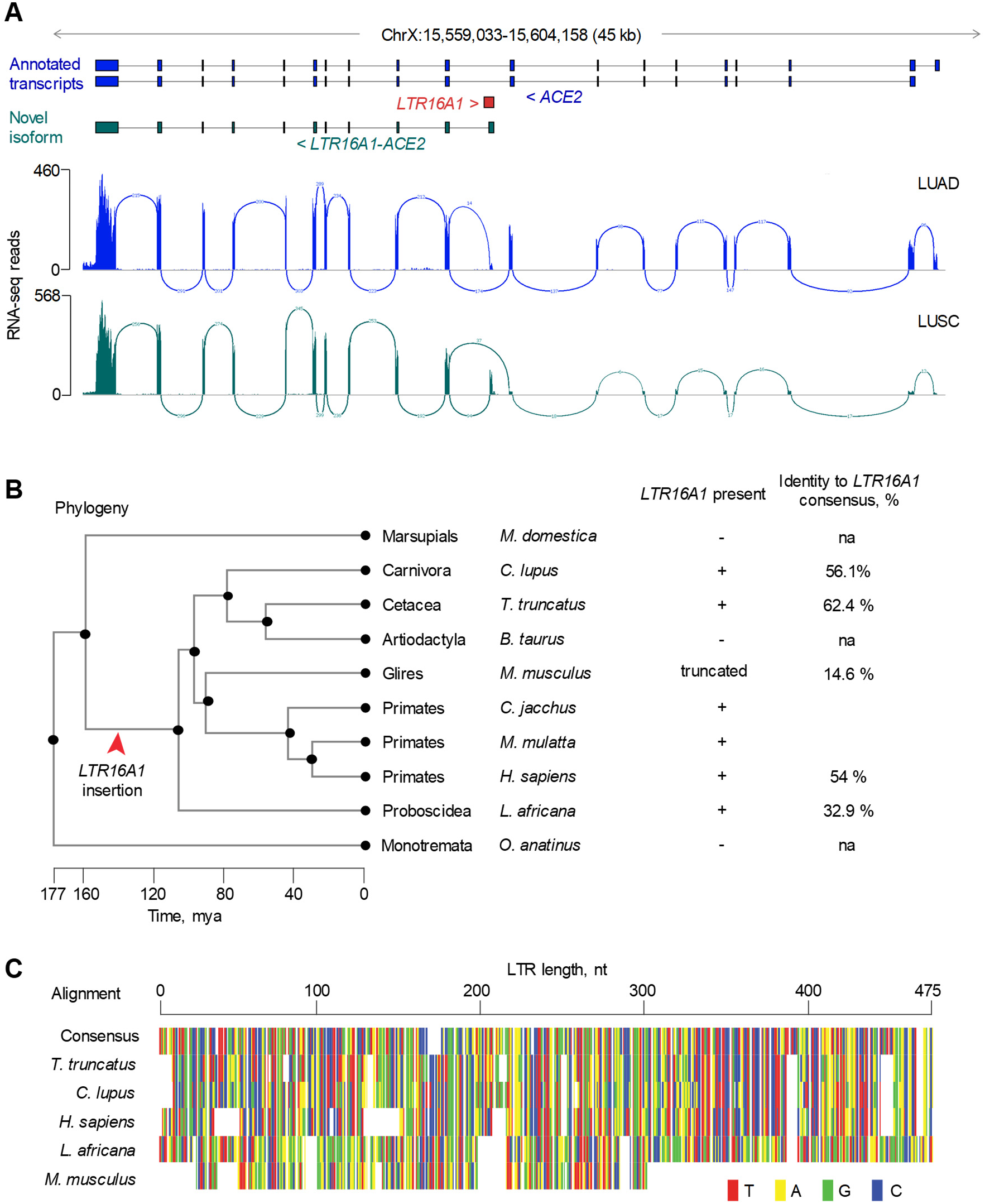
Identification of the novel *LTR16A1-ACE2* isoform. (A) GENCODE annotated transcripts at the *ACE2* locus, intronic position of the LTR16A1 element, structure of the novel *LTR16A1-ACE2* isoform and RNA-seq traces of composite LUAD and LUSC samples. Also shown is splice junction analysis of the same RNA-seq samples. (B) Phylogenetic analysis of the *LTR16A1* sequence in the indicated representative mammalian species and present sequence identity to the consensus *LTR16A1* sequence. The red arrow indicated the ancestral integration of the *LTR16A1* element. (C) Alignment of the *LTR16A1* sequence in the indicated representative mammalian species.

Phylogenetic analysis of the respective *LTR16A1* elements in the *ACE2* loci of representative mammalian species indicated that the ancestral *LTR16A1* integration predated estimated dates of mammalian radial divergence (Figure 1B). Indeed, comparative genomic analysis produced good alignment of the *LTR16A1* integration across a variety of species, with humans, dogs, and dolphins showing above 50% sequence identity to the mammalian consensus sequence of *LTR16A1* (Figure 1B, C). Of note, the *LTR16A1* integration was also present, but truncated in the mouse genome *ACE2* locus (Figure 1B, C).

The *LTR16A1-ACE2* isoform is predicted to encode a truncated *ACE2* product, retaining the last 449 amino acids of the canonical ACE2 protein and exonisation of the *LTR16A1* element creates a novel 10 amino acid sequence (MREAGWDKGG) in the putative translation product (Figure S2). Importantly, this predicted protein lacks the first 356 amino acids, including the signal peptide, substrate-binding site and domains that interact with SARS-CoV and SARS-CoV-2 spike glycoproteins (Figure S2).

To assess the relative expression of *ACE2* and *LTR16A1-ACE2* isoforms, we quantified expression of both transcripts across tissue types in the TCGA and GTEx cohorts. Consistent with recent reports (Singh et al., 2020; Ziegler et al., 2020), the full-length *ACE2* was expressed predominantly in the healthy intestine and kidney and tumours of the same histotypes (Figure S3). Expression of *LTR16A1-ACE2* followed a similar overall pattern, but with notable expression also in healthy testis, likely owing to LTR element activation as part of epigenetic reprogramming during spermatogenesis.

However, despite similar histotype distribution of *ACE2* and *LTR16A1-ACE2* expression, the ratio of the two isoforms was characteristically different between distinct histotypes and tumour types. For example, in larger TCGA patient cohorts, LUAD samples expressed higher levels of *ACE2* than of *LTR16A1-ACE2* (mean *ACE2/LTR16A1-ACE2* ratio = 5.63), whereas LUSC samples showed the opposite phenotype with higher expression of *LTR16A1-ACE2* (mean *ACE2/LTR16A1-ACE2* ratio = 0.87) (Figure 2A, B). *ACE2* and *LTR16A1-ACE2* expression and their ratios were not affected by patient gender, arguing against a strong effect of the X chromosomal location of *ACE2* on either isoform expression (Figure 2A, B). *ACE2* and *LTR16A1-ACE2* exhibited characteristic expression also within tumour types with only weak correlation between the two in the same tumour type (R^2^=0.252 for LUAD; R^2^=0.337 for LUSC), suggesting partly independent regulation.

**Figure 2.**
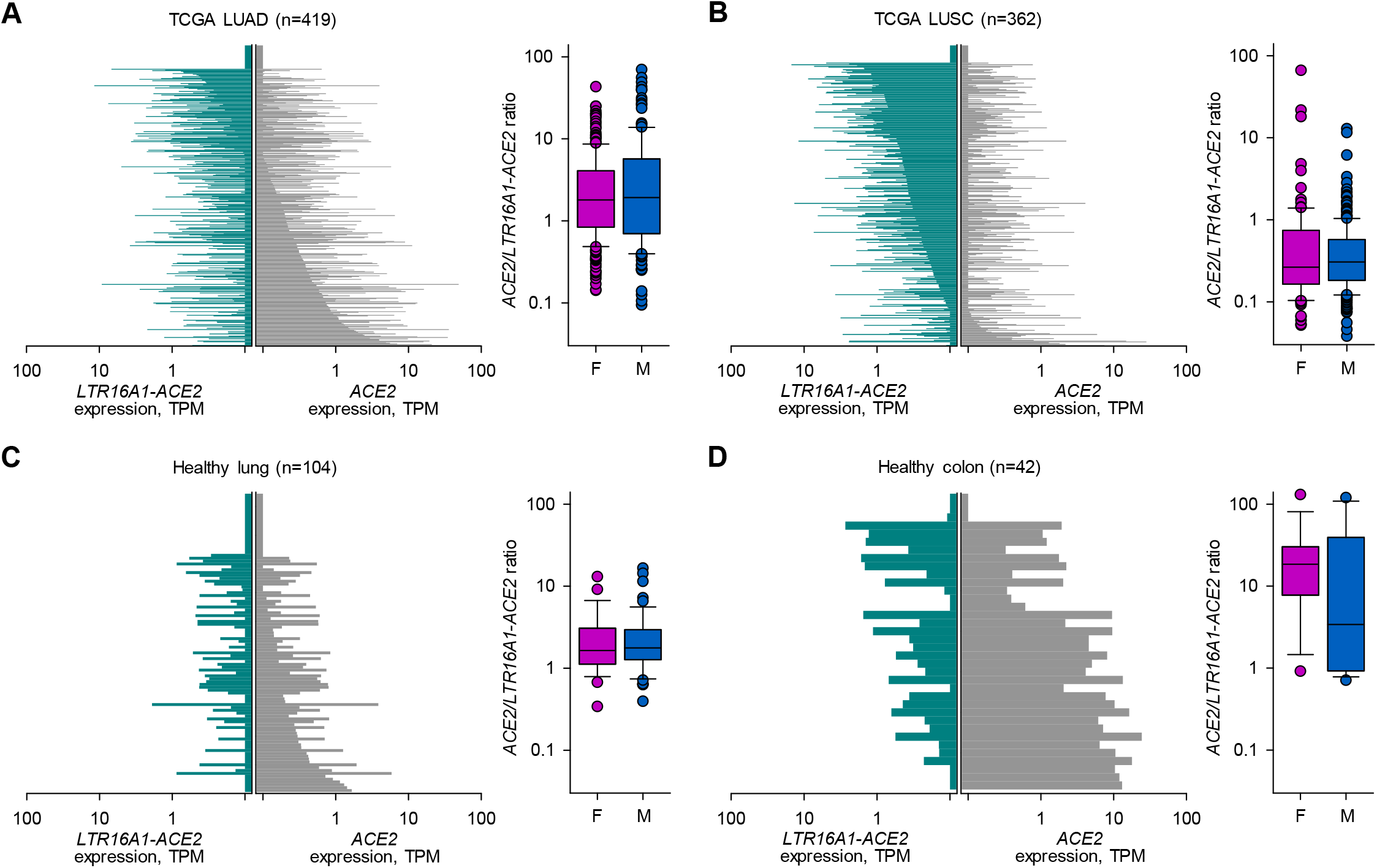
*ACE2* and *LTR16A1-ACE2* isoform expression in cancer and healthy tissues. (A) *ACE2* and *LTR16A1-ACE2* isoform expression in LUAD samples (*left*) and ratio of the two isoforms in females (F) and male (M) samples (*right*). (B) *ACE2* and *LTR16A1-ACE2* isoform expression in LUSC samples (*left*) and ratio of the two isoforms in females (F) and male (M) samples (*right*). (C) *ACE2* and *LTR16A1-ACE2* isoform expression in healthy lung samples (*left*) and ratio of the two isoforms in females (F) and male (M) samples (*right*). (D) *ACE2* and *LTR16A1-ACE2* isoform expression in healthy colon samples (*left*) and ratio of the two isoforms in females (F) and male (M) samples (*right*). In (A) to (D), each bar represents an individual samples.

In healthy lung, expression of *ACE2* and *LTR16A1-ACE2* was similar to that in LUAD, with the balance slightly in favour of the full-length form (mean *ACE2/LTR16A1-ACE2* ratio = 2.73) (Figure 2C). By contrast, healthy colon expressed considerably higher levels specifically of the full-length isoform (mean *ACE2/LTR16A1-ACE2* ratio = 26.37) (Figure 2D). These differences in *ACE2* and *LTR16A1-ACE2* expression between healthy lung and colon were again independent of gender (Figure 2C, D).

Tissue-specific patterns of *ACE2* and *LTR16A1-ACE2* expression suggested dependency on cell lineage or identity. Alternatively, they could reflect transient adaptations to the local microenvironment, such as oxygen or microbiota composition differences between lung and intestine, or even differences in cellular composition between the different compartments. To examine whether patterns of *ACE2* and *LTR16A1-ACE2* expression are linked to cell identity, we examined RNA-seq data from 933 cancer cell lines from The Cancer Cell Line Encyclopedia (CCLE). These represent homogenous cell populations, grown under standardised conditions, independently of environmental influences. Again, expression of *ACE2* and *LTR16A1-ACE2* was characteristically different between different cell lines and correlated with their anatomical origin (Figure 3A-D). Cell lines with the highest expression of *LTR16A1-ACE2* were derived from the upper aerodigestive tract, including the mouth and nose (mean *ACE2/LTR16A1-ACE2* ratio = 0.72), followed by esophageal cells lines (mean *ACE2/LTR16A1-ACE2* ratio = 1.66) and lung cell lines (mean *ACE2/LTR16A1-ACE2* ratio = 6.27). Consistent with data from primary biopsies, cells lines from the large intestine exhibited the highest expression of *ACE2*, with minimal expression of *LTR16A1-ACE2* (mean *ACE2/LTR16A1-ACE2* ratio = 16.97). The low *ACE2/LTR16A1-ACE2* ratio in the upper aerodigestive tract was highly significant when compared with other locations (p=0.0035, when compared with the lung; p=0.0023 when compared with the large intestine, Student’s *t*-test).

**Figure 3.**
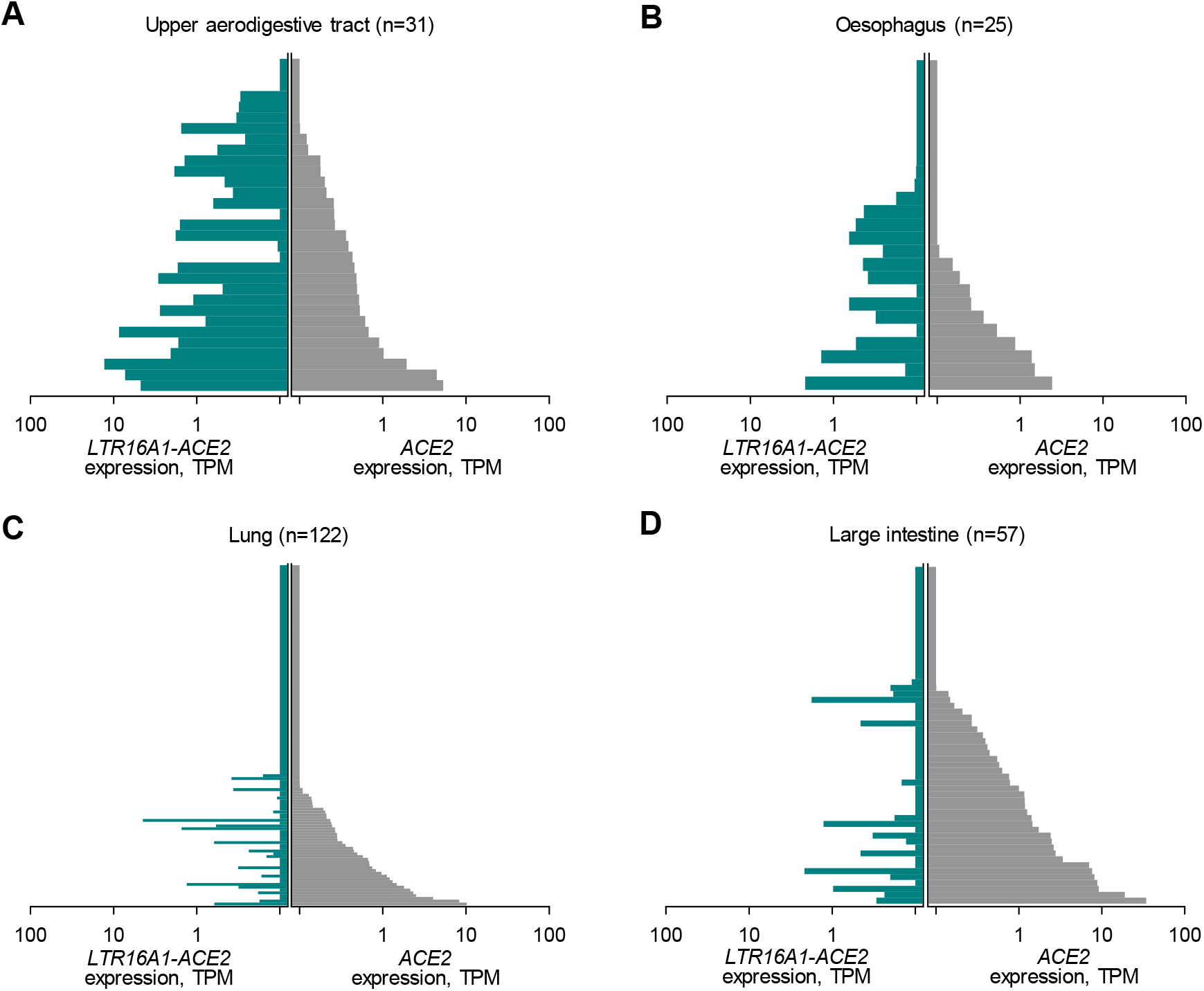
*ACE2* and *LTR16A1-ACE2* isoform expression in cell lines. (A) *ACE2* and *LTR16A1-ACE2* isoform expression in cell lines from the upper aerodigestive tract. (B) *ACE2* and *LTR16A1-ACE2* isoform expression in cell lines from the esophagus. (C) *ACE2* and *LTR16A1-ACE2* isoform expression in cell lines from the lung. (D) *ACE2* and *LTR16A1-ACE2* isoform expression in cell lines from the large intestine tract. In (A) to (D), each bar represents an individual samples.

Together, these results uncover the transcription of a novel *ACE2* isoform, initiated at an intronic *LTR16A1* retroelement, in a characteristic pattern of expression, forming a gradient from the upper aerodigestive tract (highest *LTR16A1-ACE2* expression) to the large intestine (highest *ACE2* expression).

### *LTR16A1-ACE2* and not *ACE2* transcription is IFN-responsive

*ACE2* has recently been described as a human interferon-stimulated gene (ISG), upregulated at the mRNA level following viral infection or interferon treatment (Ziegler et al., 2020). However, this conclusion was based mostly on analysis of single-cell RNA-seq data that might not have sufficient resolution to distinguish the two isoforms. Indeed, inspection of public single-cell RNA-seq data (GSE134355) (Han et al., 2020), demonstrated the limitation of such technologies, with RNA-seq reads mapping exclusively to the shared 3’ terminal exon of the *ACE2* transcripts, and therefore unable to discriminate between the isoforms (Figure S4).

To investigate the inducibility of the two isoforms by IFN or viral infection, we re-analysed public RNA-seq data (GSE147507) from normal human bronchial epithelial (NHBE) cells, treated with recombinant IFNβ or infected with SARS-CoV-2, Influenza A virus (IAV) or IAV lacking the viral NS1 protein (IAVΔNS1) (Blanco-Melo et al., 2020). None of the treatments increased expression of full-length *ACE2* (Figure 4A). In stark contrast, *LTR16A1-ACE2* expression was strongly elevated by both IAVΔNS1 infection and recombinant IFNβ treatment, compared with mock treatment (p=0.0005 and p=0.0054, respectively, Student’s *t*-test). Similar results were also obtained with analysis of lung cancer Calu-3 cells. In the absence of stimulation, Calu-3 cells express exclusively the full-length *ACE2* isoform (Figure 4B). SARS-CoV-2 infection did not affect levels of *ACE2* expression, but noticeably induced *LTR16A1-ACE2* expression (Figure 4B). Lastly, analysis of RNA-seq data from explanted lung tissue from a single COVID-19 patient demonstrated elevated expression of *LTR16A1-ACE2*, but not of *ACE2*, compared with healthy lung tissue (Figure 4C), albeit statistical comparisons were not possible in this case.

**Figure 4.**
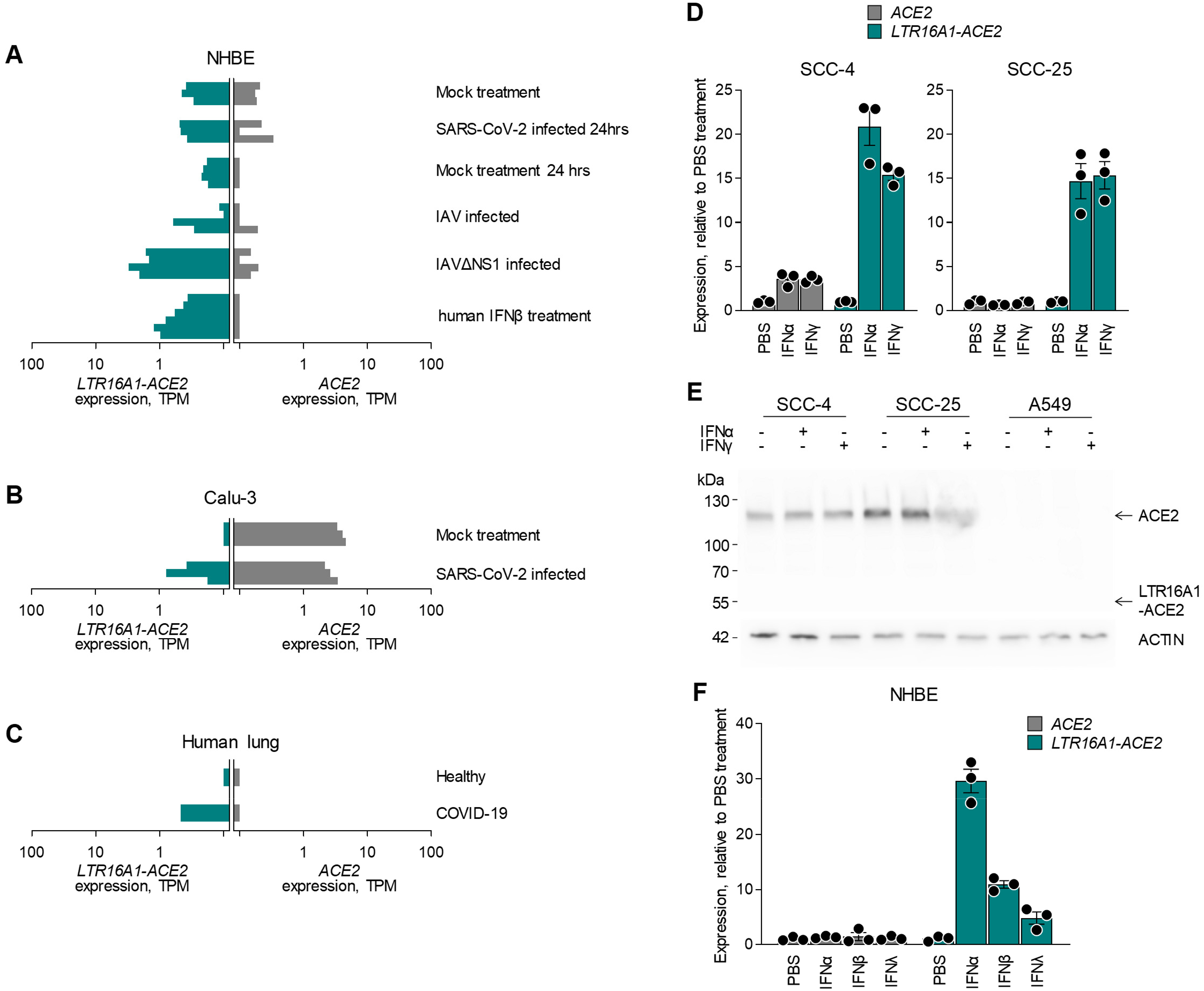
IFN inducibility of *ACE2* and *LTR16A1-ACE2* isoform expression. (A) *ACE2* and *LTR16A1-ACE2* isoform expression NHBE cells following the indicated treatment. (B) *ACE2* and *LTR16A1-ACE2* isoform expression Calu-3 cells with or without infection with SARS-CoV-2. (C) *ACE2* and *LTR16A1-ACE2* isoform expression in the lung of a COVID-19 patient and in a healthy lung. In (A) to (C), raw data were obtained form study GSE147507 and each bar represents an individual samples. (D) *ACE2* and *LTR16A1-ACE2* isoform expression, determined by RT-qPCR in SCC-4 and SCC-25 cells with or without IFN stimulation. (E) Detection of ACE2 and putative *LTR16A1-ACE2* protein product by Western blotting in cell lysates from the same cells as in (D). One representative of 2 experiments is shown. (F) *ACE2* and *LTR16A1-ACE2* isoform expression, determined by RT-qPCR in NHBE cells with or without IFN stimulation.

To further confirm the IFN-responsiveness exclusively of *LTR16A1-ACE2* expression, we used the squamous cell carcinoma cell lines SCC-4 and SCC-25, which express both isoforms. Compared with mock treatment, addition of recombinant IFNα or IFNγ had a minimal effect on *ACE2* expression in SCC-4 cells and no effect in SCC-25 cells (Figure 4D). This contrasted with very strong induction (~15-fold) of *LTR16A1-ACE2* expression by either type of IFN in both cell lines (Figure 4D). Lack of *ACE* responsiveness to IFN stimulation was additionally confirmed at the protein level, where neither IFNα nor IFNγ affected levels of full-length ACE2, detected by Western blotting in SCC-4 and SCC-25 cells or in A549 cells, which express neither isoform and were used as a negative control (Figure 4E). Of note, despite strong upregulation at the mRNA level and despite using an antibody targeting the C-terminus of ACE2 present in both protein products, we were unable to detect a truncated form that would correspond to the *LTR16A1-ACE2* translation product (Figure 4E).

To confirm the differential IFN inducibility of *ACE2* and *LTR16A1-ACE2* expression, we stimulated NHBE cells with IFNα, IFNβ or IFNλ, as previously described (Major et al., 2020). Again, treatment with none of the IFNs had any measurable effect on *ACE2* expression in these primary cells (Figure 4F). This contrasted with robust induction of *LTR16A1-ACE2* expression, particularly by IFNα (Figure 4F).

Collectively, these data demonstrate that type I, II and III IFNs stimulate transcription of the *ACE2* isoform driven by the alternative *LTR16A1*, but not the canonical *ACE2* promoter.

### The *LTR16A1-ACE2* protein product is not stable

Splicing from the *LTR16A1* promoter to exon 10 of *ACE2* is in-frame and therefore the last 449 amino acids are shared between ACE2 protein and the putative LTR16A1-ACE2 protein. Nevertheless, we have been unable to detect the latter in cells naturally expressing the *LTR16A1-ACE2* transcript (Figure 4E). To explore the protein-coding potential of the *LTR16A1-ACE2* transcript, we cloned the coding sequences of both isoforms into the pcDNA3.1 mammalian expression vector and transfected HEK293T cells. While *ACE2*-transfected HEK293T cells produced readily detectable ACE2 protein, no protein of the predicted size was detectable in *LTR16A1-ACE2*-transfected cells (Figure 5A). We quantified levels of enzymatically active ACE2 and found, as expected, strong enzymatic activity in lysates from *ACE2*-transfected, but not *LTR16A1-ACE2*-transfected cells (Figure 5B).

**Figure 5.**
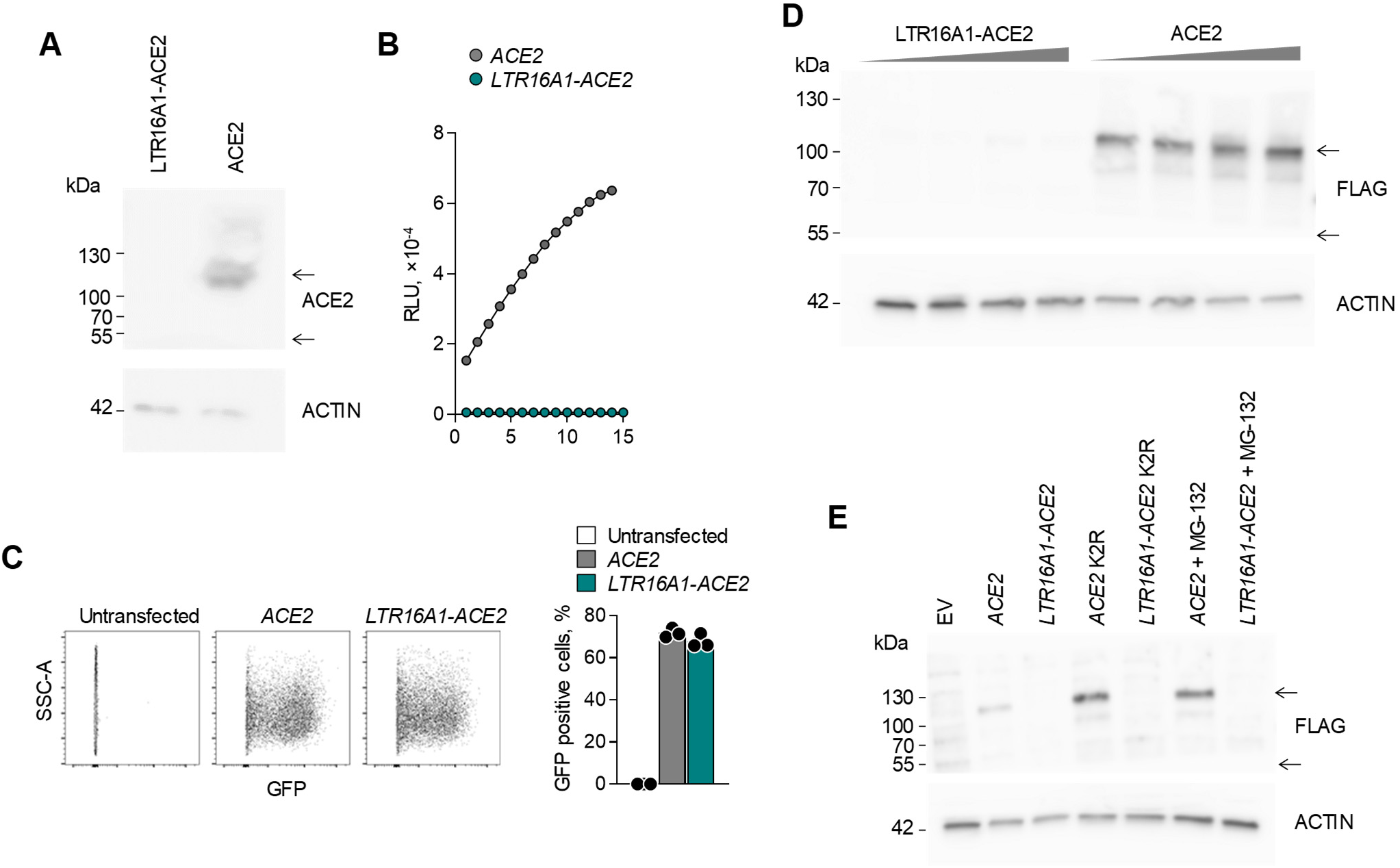
Stability of the *ACE2* and *LTR16A1-ACE2* translation products. (A) Detection of ACE2 and putative *LTR16A1-ACE2* protein product by Western blotting in cell lysates from HEK293T cells transfected to express either isoform. One representative of 4 experiments is shown. (B) ACE2 enzymatic activity in the supernatant of the same cells as in (A). (C) Flow cytometric detection of GFP expression (*left*) and quantitation of GFP-expressing cells (*right*) in HEK293T cells transfected to express either isoform in conjunction with a FLAG tag and GFP, linked by a P2A peptide. One representative of 2 experiments is shown. (D) Detection of ACE2 and putative *LTR16A1-ACE2* protein product by Western blotting for the FLAG tag in cell lysates from the same cells as in (C). Titration the transfection plasmids used is also indicated. One representative of 2 experiments is shown. (E) Detection of ACE2 and putative *LTR16A1-ACE2* protein product by Western blotting for the FLAG tag in cell lysates from HEK293T cells transfected to express either wild-type isoform or either isoform with the two lysine residues mutated (K2R) (all in conjunction with a FLAG tag and GFP, linked by a P2A peptide). HEK293T cells transfected to express the wild-type isoforms were treated with the MG-132 inhibitor. One representative of 2 experiments is shown.

To confirm the lack of protein production from *LTR16A1-ACE2* transcript, we cloned the coding sequences of both isoforms into the pcDNA3.1-DYK-P2A-GFP expression vector, which adds both a FLAG tag and GFP as part of the protein product. Expression of GFP was comparable in *ACE2*-transfected and *LTR16A1-ACE2*-transfected cells, suggesting that the single RNA molecule that encodes for both the *LTR16A1-ACE2* product and GFP is stable and translated (Figure 5C). Despite that, while full-length ACE2 was detectable across a range of plasmid concentrations, we could not detect the predicted *LTR16A1-ACE2* protein product with antibodies to the FLAG tag (Figure 5D).

Lysine residues 625 and 702 in the full-length ACE2 protein have been described to be ubiquitinated and may contribute to its proteosomal degradation (Stukalov et al., 2020). We generated a K625R K702R (K2R) mutant of full-length ACE2, which increased protein levels, compared to the wild-type ACE2 (Figure 5E). We have introduced the same mutations in the corresponding residues of the predicted *LTR16A1-ACE2* protein product, K279R K356R, which were similarly accessible for ubiquitination (Figure S5). However, we were unable to detect stable protein following transfection with the *LTR16A1-ACE2* K2R mutant (Figure 5E). Consistent with this, the addition of the proteasome inhibitor MG-132 was sufficient to increase protein levels of ACE2, but did not rescue the *LTR16A1-ACE2* protein product (Figure 5E). These data suggest that the latter protein is subject to post-translational degradation through a proteasome-independent mechanism and therefore unlikely to exert significant biological activity.

## Discussion

Regulation of ACE2 expression and function is critical both in physiology and pathology (Hamming et al., 2007). The use of ACE2 as a primary receptor for entry by pandemic coronaviruses, SARS-CoV and SARS-CoV-2 highlighted the potential effect of changes in ACE2 expression, particularly in response to IFN, on the course or severity of COVID-19 (Ziegler et al., 2020). Here, we showed that *ACE2* transcription and protein production is not responsive to IFN. Instead, we describe a novel RNA isoform, *LTR16A1-ACE2*, that is highly responsive to IFN stimulation, but encodes a truncated and unstable protein product. This isoform showed distinct patterns of expression along the aerodigestive and gastrointestinal tracts and was likely responsible for the apparent IFN inducibility of *ACE2* expression reported by analysis of single-cell RNA-seq data (Ziegler et al., 2020). We further show that transcription of this novel isoform is initiated by an intronic LTR retroelement, which functions as a cryptic, IFN-responsive promoter, adding further evidence for the widespread involvement of such retroelements in gene regulatory networks.

Indeed, endogenous retroelements comprise nearly half the human genome and can affect many host processes (Burns and Boeke, 2012; Feschotte and Gilbert, 2012; Kassiotis and Stoye, 2016). LTR retroelements represent an abundant source of promoters, enhancers, and polyadenylation sequences that can modulate the expression and structure of neighbouring genes (Thompson et al., 2016), as with *ACE2*. For instance, retroelements serve as promoters or enhancers for a number of ISGs, conferring IFN inducibility, exemplified in the case of *AIM2* (Chuong et al., 2016). Retroelements may further modify the function of ISGs and we have recently described a novel isoform of the ISG *CD274* (encoding PD-L1) that produces a truncated form through retroelement exonisation (Ng et al., 2019).

The use of the intronic *LTR16A1* element as the promoter for the *LTR16A1-ACE2* isoform explains its independent regulation from that of the full-length *ACE2* isoform. In addition to IFN inducibility, the cryptic *LTR16A1* promoter also confers tissue-specific expression, with the highest levels seen in the upper aerodigestive tract, where it can be the predominant isoform.

In contrast, the canonical *ACE2* isoform far exceeds expression of the *LTR16A1-ACE2* isoform in the lower gastrointestinal tract. It is theoretically possible that the balance of *LTR16A1-ACE2* and full-length *ACE2* isoforms plays a role in the spread of SARS-CoV-2, particularly in the upper aerodigestive tract, or that RNA or protein products of *LTR16A1-ACE2* are involved in other pathological or physiological processes. However, the lack of a stable protein corresponding to the *LTR16A1-ACE2* coding sequence argues that this is unlikely.

Independently of any functional significance, expression of the *LTR16A1-ACE2* isoform needs to be carefully considered in studies examining *ACE2* regulation at the transcriptional level (Singh et al., 2020; Ziegler et al., 2020). The description of this novel isoform highlights the need to validate single-cell RNA-seq data with orthogonal approaches. While single-cell RNA-seq initiatives are an invaluable resource and allow for rapid identification of cell types that express a gene of interest, coverage and read depth are largely insufficient to distinguish between isoforms. Technological advances to improve sequencing depth and bioinformatic tools to impute missing values are rapidly progressing; in the meantime, long-read sequencing techniques to quantify transcript isoforms and confirmation of protein expression levels can be incorporated into existing workflows.

This work established *LTR16A1-ACE2* as the predominant form of *ACE2* following viral infection or recombinant interferon treatment, including in the SARS-CoV-2-infected lung. The suggestion that *ACE2* is an ISG raised fears that therapeutic interferon could be detrimental (Ziegler et al., 2020); however, we find that full-length ACE2 is not increased at the mRNA or protein level. Under none of the conditions tested, did *LTR16A1-ACE2* produce any detectable protein. However, we cannot rule out the possibility that it may produce a protein product or fragments thereof under certain conditions *in vivo*. Nevertheless, it is worth noting that the predicted *LTR16A1-ACE2* protein product does not contain the residues required for SARS-CoV-2 spike glycoprotein binding (Shang et al., 2020) and is thus unlikely to contribute to viral spread. These results reconcile the apparent discrepancy between the interferon inducibility of *ACE2* with promising data showing improved outcomes in COVID-19 following interferon treatment (Hung et al., 2020; Wang et al., 2020).

## Methods

### KEY RESOURCES TABLE

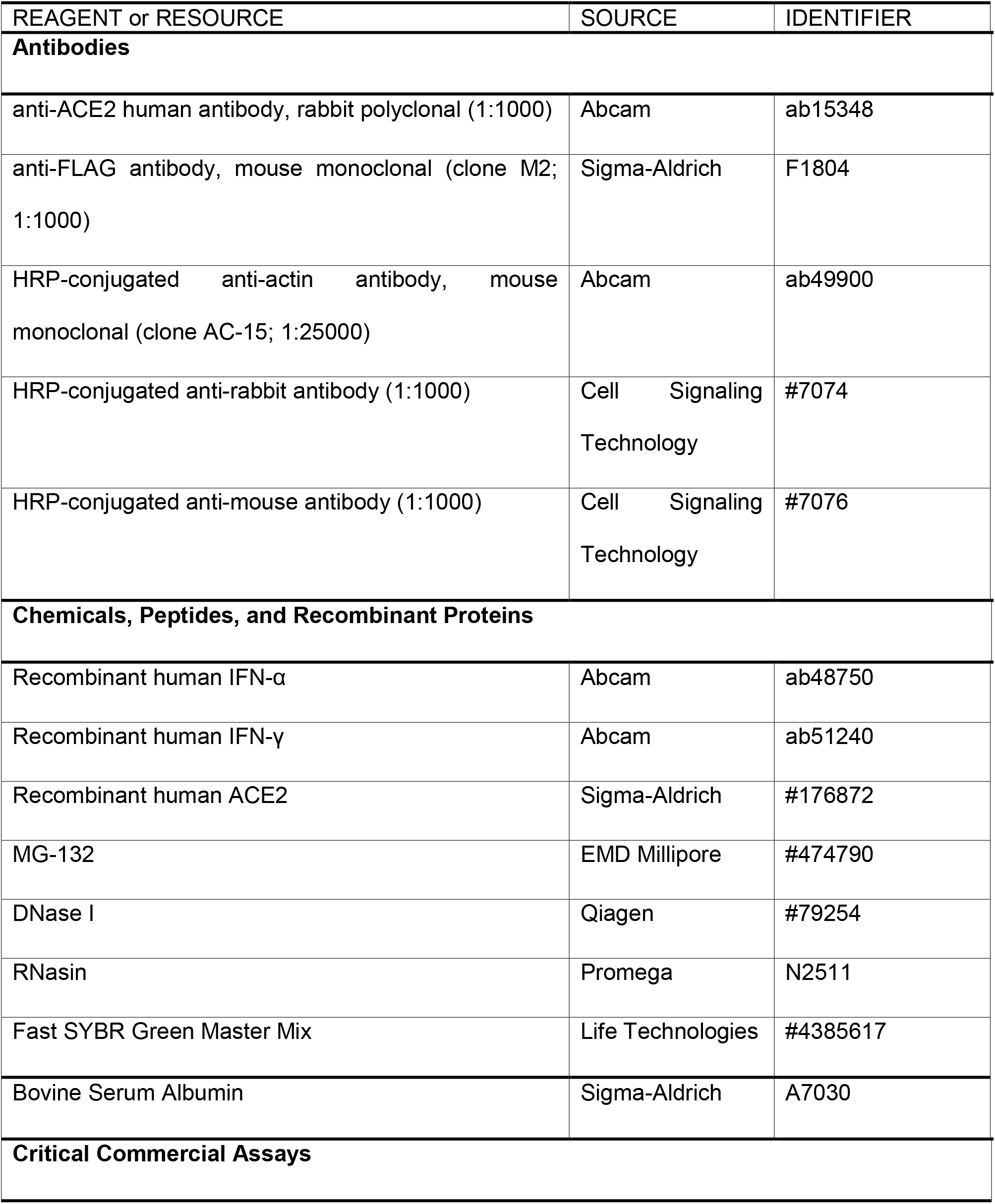
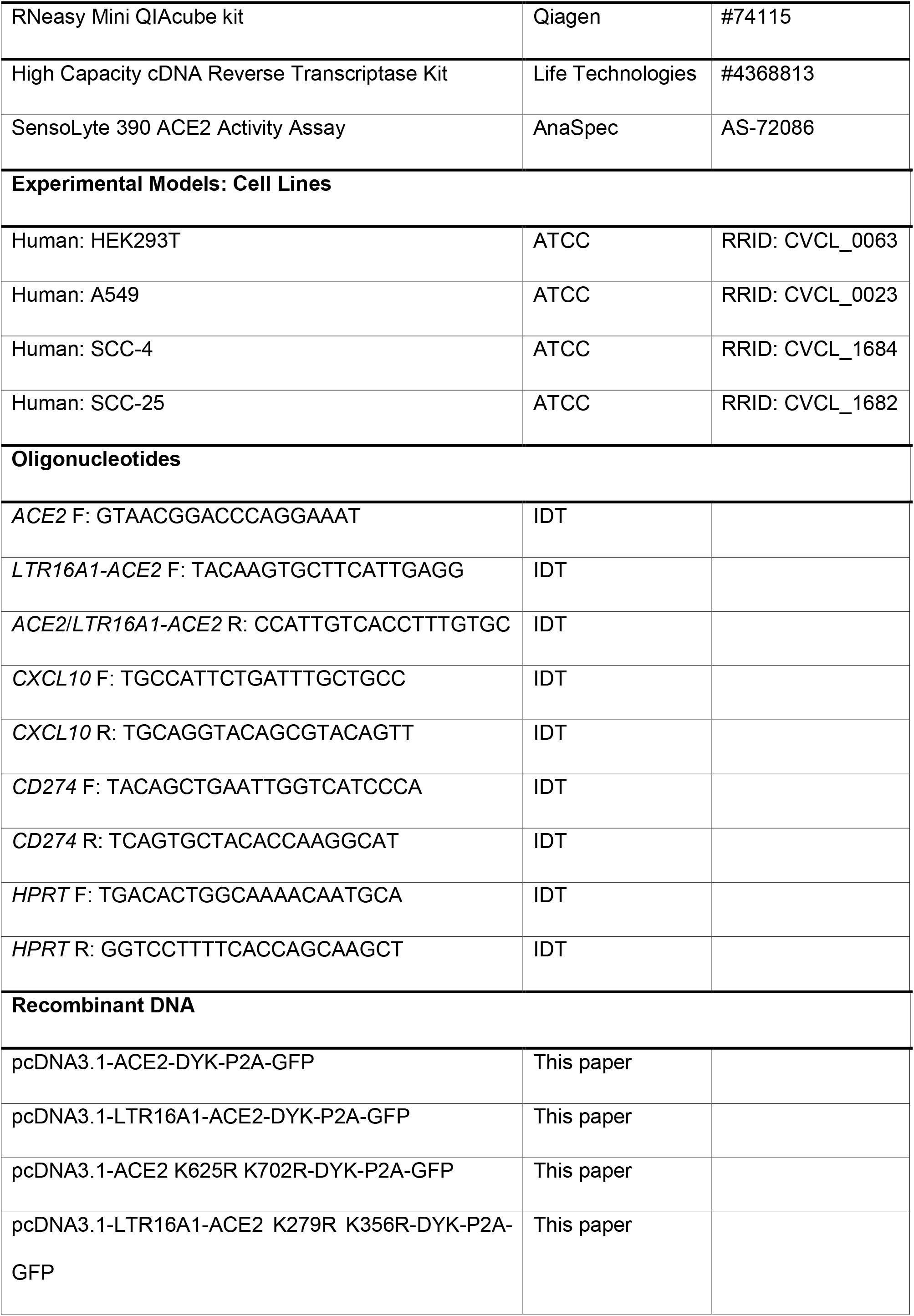
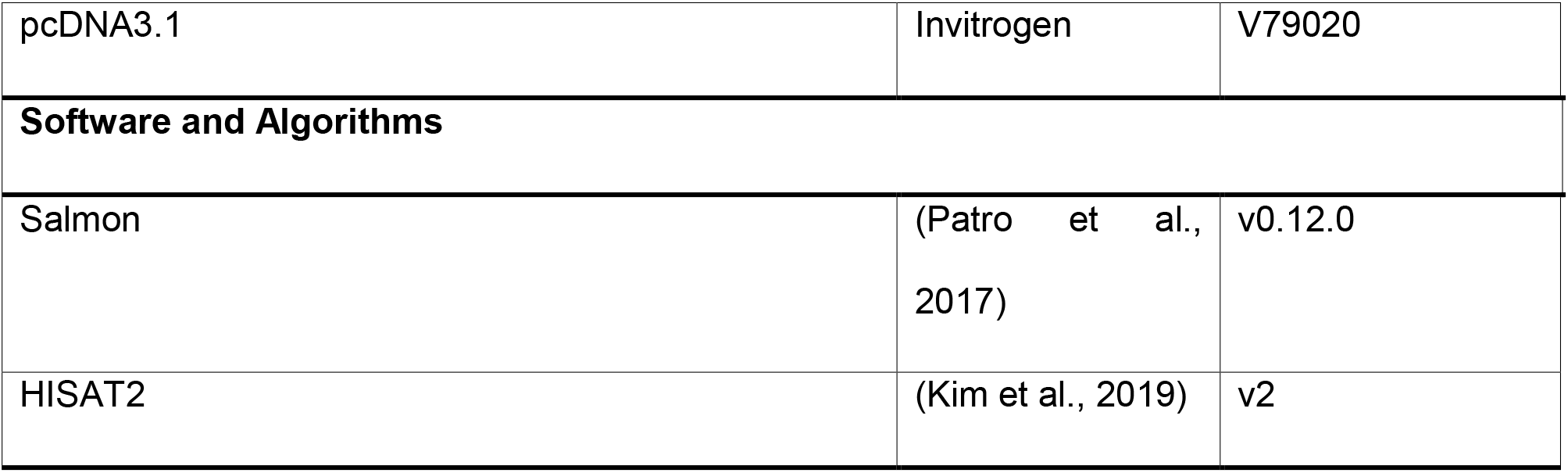

## RESOURCE AVAILABILITY

### Lead Contact

Further information and requests for resources and reagents should be directed to and will be fulfilled by the Lead Contact, George Kassiotis

### Materials Availability

DNA constructs and other research reagents generated by the authors will be distributed upon request to other research investigators.

### Data and Code Availability

The published article includes all data generated or analysed during this study, and are summarized in the accompanying figures and Supplemental materials.

## EXPERIMENTAL MODEL AND SUBJECT DETAILS

### Cell lines

HEK293T, A549, SCC-4, and SCC-25 cells were obtained from and verified as mycoplasma free by the Cell Services facility at the Francis Crick Institute. All cell lines were validated by DNA fingerprinting. HEK293T and A549 cells were grown in Iscove’s Modified Dulbecco’s Medium (Sigma-Aldrich) supplemented with 5% fetal bovine serum (Thermo Fisher Scientific), L-glutamine (2 mmol/L, Thermo Fisher Scientific), penicillin (100 U/mL, Thermo Fisher Scientific), and streptomycin (0.1 mg/mL, Thermo Fisher Scientific). SCC-4 and SCC-25 cells were grown in Dulbecco’s Modified Eagle Medium/Nutrient Mixture F-12 (Gibco) supplemented with 10% fetal bovine serum (Thermo Fisher Scientific), L-glutamine (2 mmol/L, Thermo Fisher Scientific), penicillin (100 U/mL, Thermo Fisher Scientific), and streptomycin (0.1 mg/mL, Thermo Fisher Scientific). NHBE cells were cultured as previously described (Major et al., 2020)

## METHOD DETAILS

### Transcript identification and read mapping

Transcripts were previously assembled on a subset of the RNAseq data from The Cancer Genome Atlas (TCGA) (Attig et al., 2019). The alternative promoter within *ACE2* was more highly expressed in lung squamous carcinomas than the canonical isoform, prompting us to investigate its biology. RNAseq data from TCGA, GTEx, CCLE, and other studies were aligned to the cancer-tissue transcriptome assembly using GNU parallel (Tange, 2011) and Salmon v0.12.0 (Patro et al., 2017). Splice junctions were visualised using the Integrative Genome Viewer v2.4.19 (Thorvaldsdóttir et al., 2013).

### LTR16A1 Sequence alignments

To identify the integration time of *LTR16A1* into the *ACE2* locus, we first compared the Homo sapiens *LTR16A1* to the *LTR16A1* consensus sequence in repBase. Based on 285 identical bases across its 396 bp length, the insertion is expected to be ~127 million years old based on the human neutral substitution rate at 2.2 × 10^−9^ substitutions per site per year. To find evidence for insertion of the *LTR16A1* site before the split of the major mammalian lineages, we used the UCSC liftover utility to find the *ACE2* gene locus in Rhesus macaque (rheMac10 assembly), marmoset (caljac3 assembly), mouse (mm10 assembly), dog (canFam3 assembly), african elephant (loxAfr3 assembly), bottle-nose dolphin (Turtru2 assembly), cow (bosTau9 assembly), opossum (monDom5 assembly) and platypus (ornAna2). We used the MUSCLE aligner on default settings to build a global alignment of human to rhesus macaque and marmoset, and then aligned all other species to the profile, reverting the strand of the whole sequence for mouse, elephant, cow and opossum due to whole gene inversions. We then used muscle *-refine* on overlapping 30,000 column blocks to refine the alignment locally. The illustration of the lineage tree including node times are taken from timetree.org.

### Analysis of bulk RNA-seq and single-cell RNA-seq data

Bulk RNA-seq data were downloaded from study GSE147507 (Blanco-Melo et al., 2020). Reads were adapter trimmed and filtered for minimal 35nt sequences using Trimmomatic. Since some samples were infected with SARS-CoV-2 *in vitro*, we identified and removed viral reads using BowTie2 (*seedlength* 30nt) to align reads to the Wuhan region reference genome (MN908947). Subsequently, reads were mapped with HISAT2 (optional parameters *−p 8 −q −k 5*) against GRCh38 reference chromosome assembly and transcripts were quantified against our custom transcriptome assembly (Attig et al., 2019) using Salmon (v0.12.0) (Patro et al., 2017).

For single-cell RNA-seq data analysis, we downloaded the raw paired end sequencing reads as unmapped bam files from study GSE134355 (Han et al., 2020), which were already demultiplexed, with one individual per tissue per sample. We then used the *DropSeq*:*picard* toolbox (v2.3.0) to recapitulate processing of HCL samples as documented on ‘https://github.com/ggjlab/HCL’. In summary, this includes trimming polyA ends from each primary RNA sequencing read and tagging it with the cellular and molecular adapter sequence contained in the secondary read (BASE_RANGE=1-6:22-27:43-48 and BASE_RANGE=49-54, respectively). All reads were then mapped with HISAT2 (optional parameters *−p 8 −q −k 5*) against GRCh38 reference chromosome assembly. The HISAT2 index here was built with the *--exon* / *--ss* option to cover all known splice sites annotated in the GENCODE v34 *basic* annotation. The cellular and molecular barcode sequences were recovered using the *MergeBamAlignment* utility in *picard*.

### Expression vectors

Open reading frames encoding *ACE2*, *LTR16A1-ACE2*, and respective lysine mutants were synthesized and cloned into the pcDNA3.1-DYK-P2A-GFP mammalian expression vector. Gene synthesis, cloning, and mutagenesis were performed by GenScript and verified by sequencing. Cells were transfected using GeneJuice (EMD Millipore) and harvested 48 hrs post-transfection for downstream assays.

### Cell stimulation

For interferon stimulation experiments, 2 × 10^5^ SCC-4 and SCC-25 cells were stimulated with 100 ng/mL IFN-α or IFN-γ (Abcam) or PBS for 48 hrs. For proteasome inhibition experiments, cells were cultured in 20 μM MG-132 (EMD Millipore) 24 hrs after transfection and harvested 48 hrs after transfection. NHBE cells were stimulated for 4 hrs with 1000 ng/ml IFNα, 100ng/ml IFNβ or 100ng/ml IFNλ were used in a previous study (Major et al., 2020), and stored cDNA was analysed by RT-qPCR in this study.

### Western blot

Cell lysates in RIPA buffer were resuspended in SDS buffer, heat denatured at 95°C for 10 min, run on a 4-20% gel (Biorad), transferred to a PVDF membrane (Biorad), and blocked in 5% (w/v) bovine serum albumin fraction V (Sigma-Aldrich) in TBS-T. Membranes were incubated with primary antibodies to ACE2 (1:1000; Abcam), FLAG (1:1000, clone M2; Sigma-Aldrich), HRP-conjugated secondary antibodies to rabbit and mouse (1:1000; Cell Signaling Technology), and HRP-conjugated actin (1:25000; Abcam). Blots were visualized by chemiluminescence on an Amersham Imager 600 (GE Healthcare).

### Reverse transcriptase-based quantitative PCR (RT-qPCR)

Total RNA from cell lines was isolated using the QIAcube (Qiagen), and cDNA synthesis was carried out with the High Capacity Reverse Transcription Kit (Applied Biosystems) with an added RNase inhibitor (Promega). Purified cDNA was used to quantify *ACE2* and *LTR16A1-ACE2* using variant-specific primers.

The variant-specific forward primers were:

*ACE2* | F: GTAACGGACCCAGGAAAT
*LTR16A1-ACE2* | F: TACAAGTGCTTCATTGAGG

For both variants, the following common reverse primer was used:

Common primer | R: CCATTGTCACCTTTGTGC

Interferon-inducible genes were amplified using the following primers:

*CXCL10* | F: TGCCATTCTGATTTGCTGCC R: TGCAGGTACAGCGTACAGTT
*CD274* | F: TACAGCTGAATTGGTCATCCCA R: TCAGTGCTACACCAAGGCAT

For amplification of a conserved house-keeping gene, the following *HPRT*-specific primers were used:

*HPRT* | F: TGACACTGGCAAAACAATGCA; R: GGTCCTTTTCACCAGCAAGCT

Values were normalised to *HPRT* expression using the ΔC_T_ method.

### Enzyme assays

ACE2 activity in cell lysates was measured using the SensoLyte 390 ACE2 Activity Assay (AnaSpec) according to manufacturer’s instructions. Recombinant human ACE2 (Sigma-Aldrich) was used as a positive control.

### Flow Cytometry

For GFP detection, single-cell suspensions were run on a LSR Fortessa (BD Biosciences) running BD FACSDiva v8.0 and analysed with FlowJo v10 (Tree Star Inc.) analysis software.

## QUANTIFICATION AND STATISTICAL ANALYSIS

Statistical comparisons were made using GraphPad Prism 7 (GraphPad Software) or SigmaPlot 14.0. Parametric comparisons of normally distributed values that satisfied the variance criteria were made by unpaired Student’s *t*-tests or One Way Analysis of variance (ANOVA) tests. Data that did not pass the variance test were compared with non-parametric two-tailed Mann–Whitney Rank Sum tests or ANOVA on Ranks tests.

## Acknowledgments

We are grateful for assistance from the Scientific Computing and Flow Cytometry facilities at the Francis Crick Institute. The results shown here are in whole or part based upon data generated by The Cancer Genome Atlas (TCGA) Research Network (http://cancergenome.nih.gov). The Genotype-Tissue Expression (GTEx) Project was supported by the Common Fund of the Office of the Director of the National Institutes of Health, and by NCI, NHGRI, NHLBI, NIDA, NIMH, and NINDS. This work benefited from data assembled by the CCLE consortium. This work was supported by the Francis Crick Institute (FC001099, FC001206), which receives its core funding from Cancer Research UK, the UK Medical Research Council, and the Wellcome Trust; and by the Wellcome Trust (102898/B/13/Z).

---

**Figure S1.**
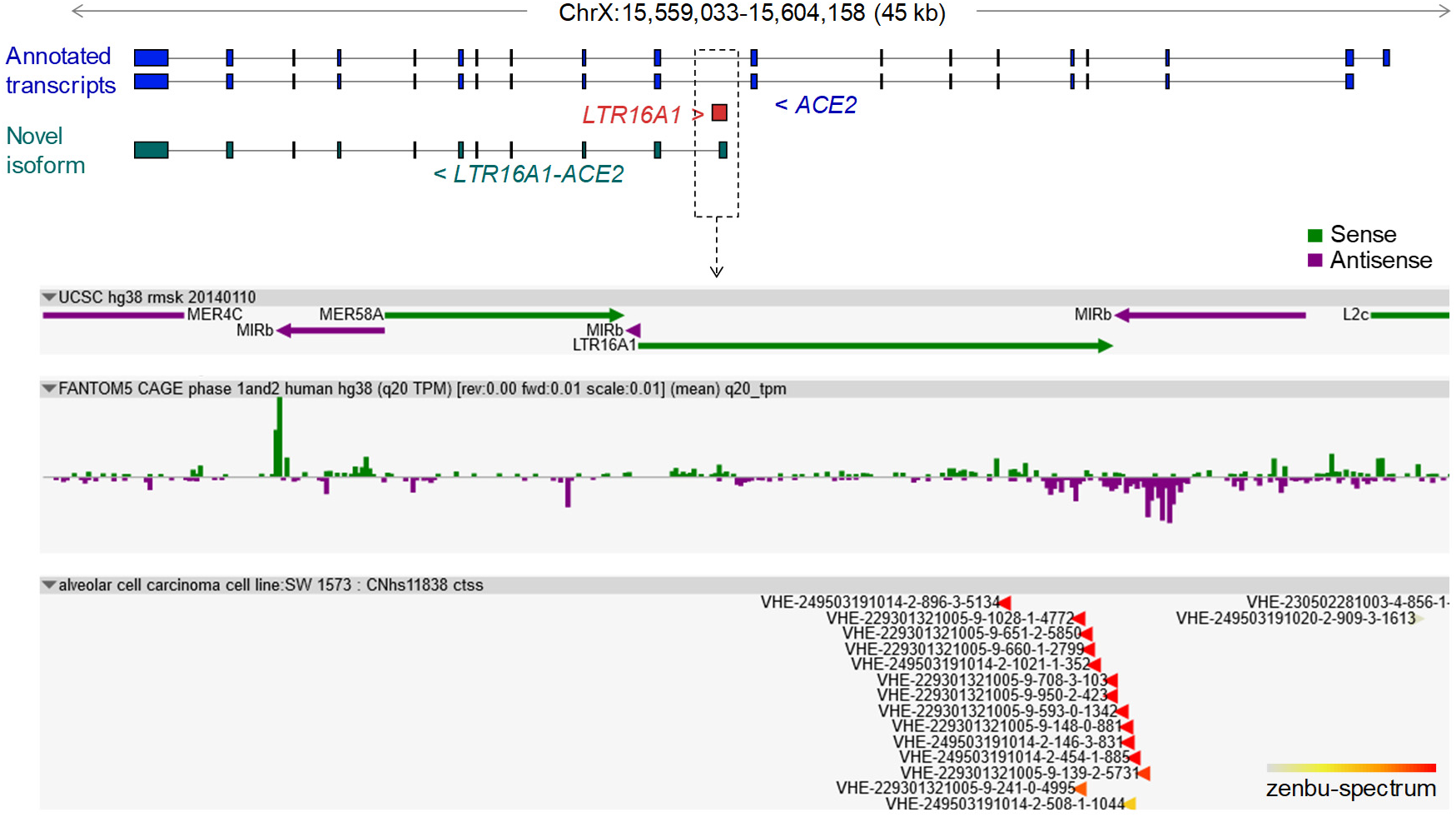
CAGE support for transcriptional initiation of the *LTR16A1-ACE2* transcript. Normalized data from the FANTOM Consortium and the RIKEN PMI and CLST (DGT) for transcription start sites in the proximity of the intronic *LTR16A1* element in the *ACE2* locus. Both the sense and antisense orientations are depicted. Data were visualized with the *zenbu* online viewer (https://fantom.gsc.riken.jp/zenbu) for FANTOM5 Human hg38 promoterome.

**Figure S2.**
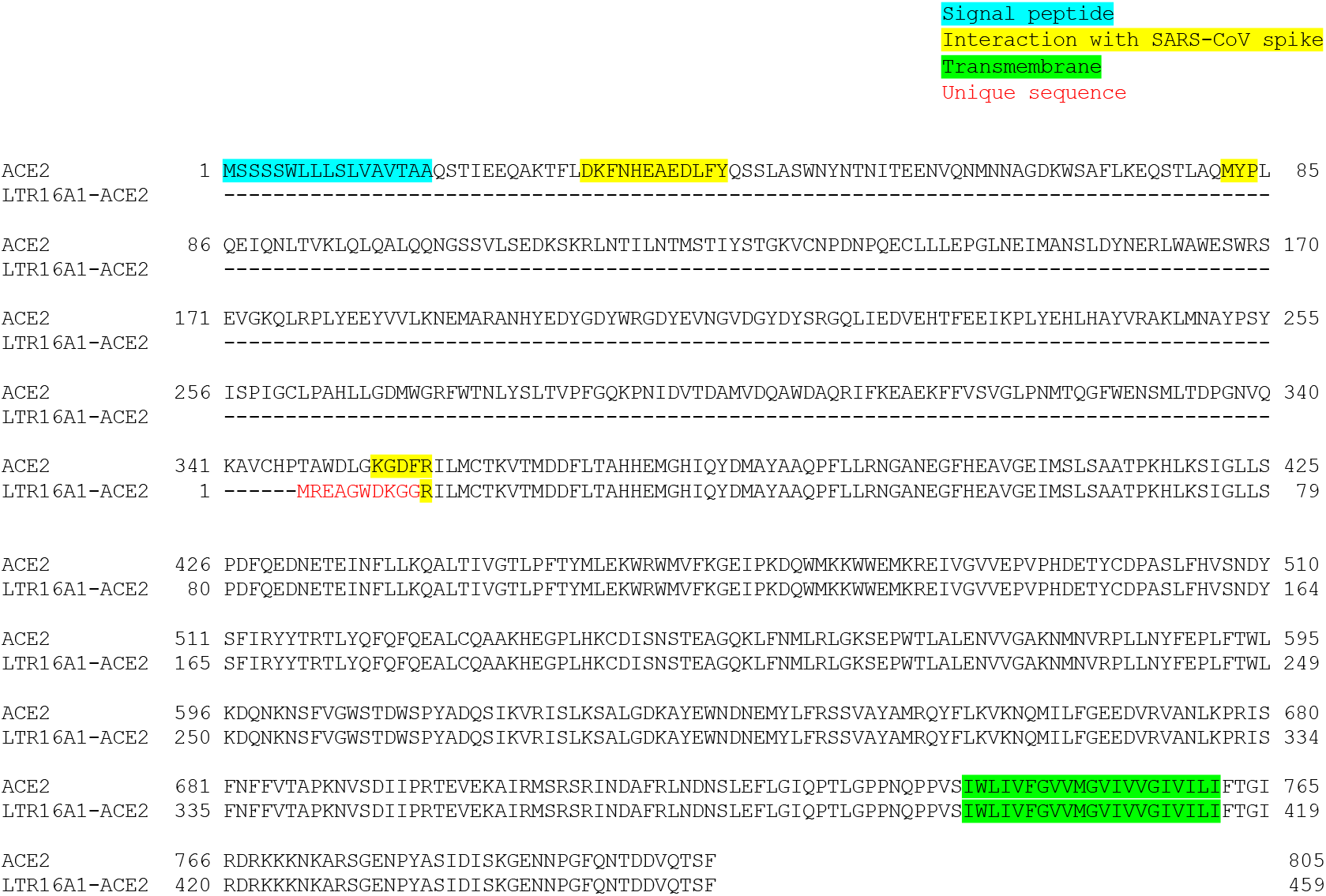
Protein sequence alignment of ACE2 and the predicted *LTR16A1-ACE2* product. The predicted *LTR16A1-ACE2* translation product is a 459-amino acid protein lacking the indicted single peptide, domains interacting with SARS-CoV spike glycoprotein, but retaining the transmembrane domain. The novel 10-amino acid sequence created by *LTR16A1* exonisation is also shown.

**Figure S3.**
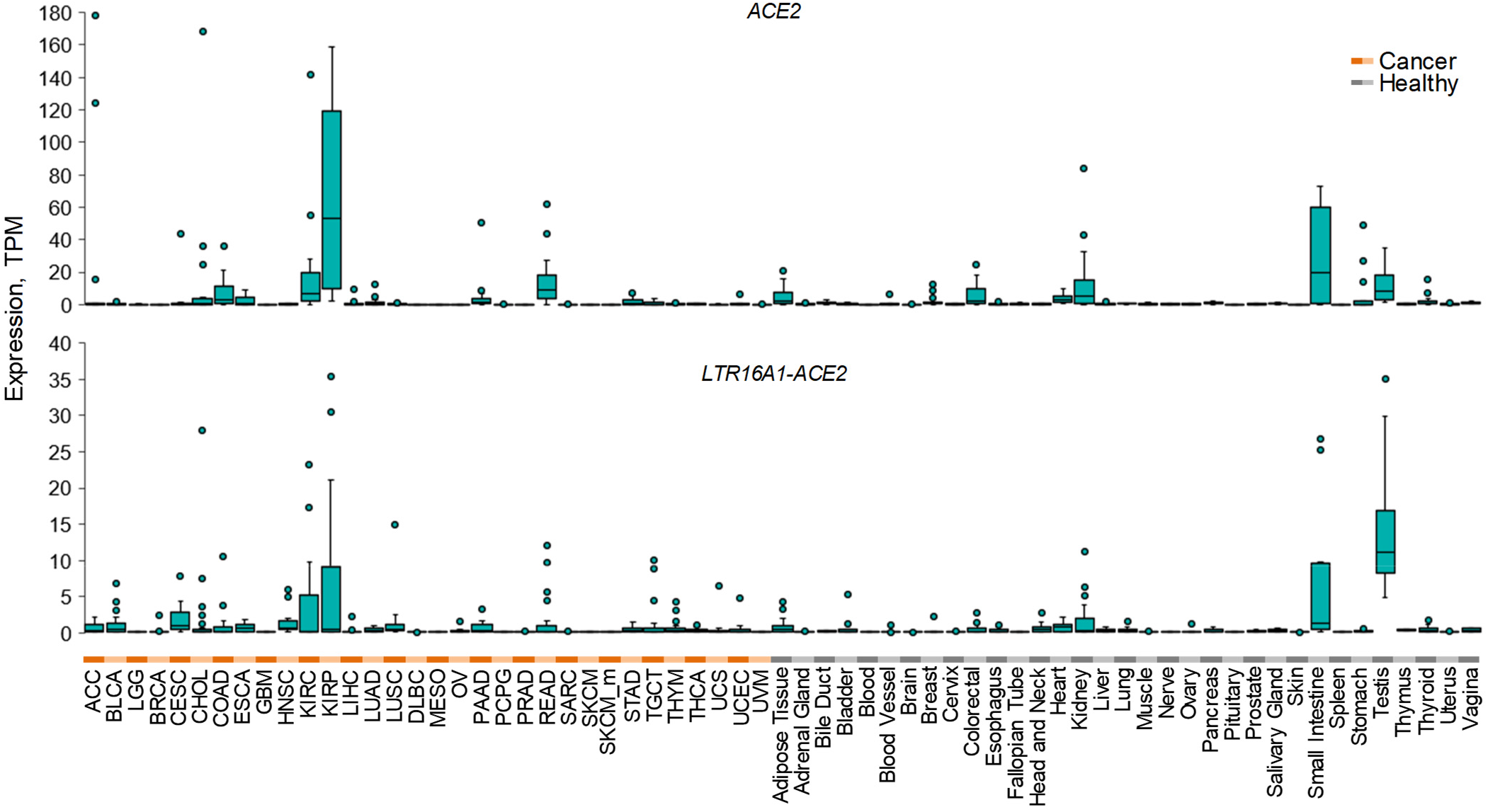
Expression of *ACE2* and *LTR16A1-ACE2* isoforms in cancer and healthy tissues. Box plots of *ACE2* and *LTR16A1-ACE2* isoforms expression in cancer patient and healthy control samples from TCGA and GTEx. For each cancer type, 24 samples were included (a total 768 samples), whereas for respective healthy tissues a total of 813 samples were included, varying between 2 and 156 per tissue type.

**Figure S4.**
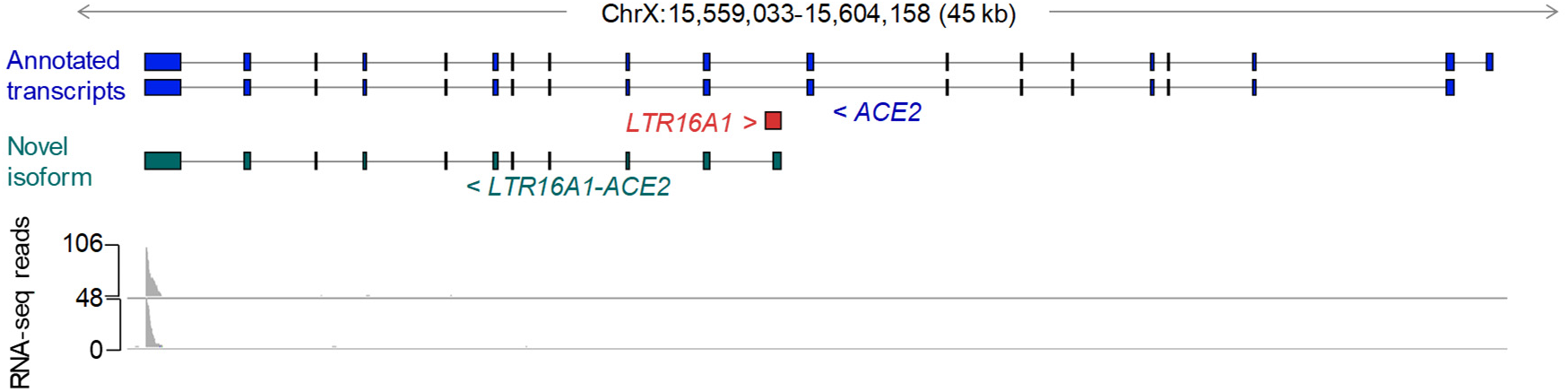
Single-cell RNA-seq coverage of the ACE2 locus. RNA-seq trace of two multiplexed samples from adult lung, obtained from study GSE134355. Note the lack of coverage across the entire locus with the exception of only the 3’ end of the last exon, shared between the isoforms.

**Figure S5.**
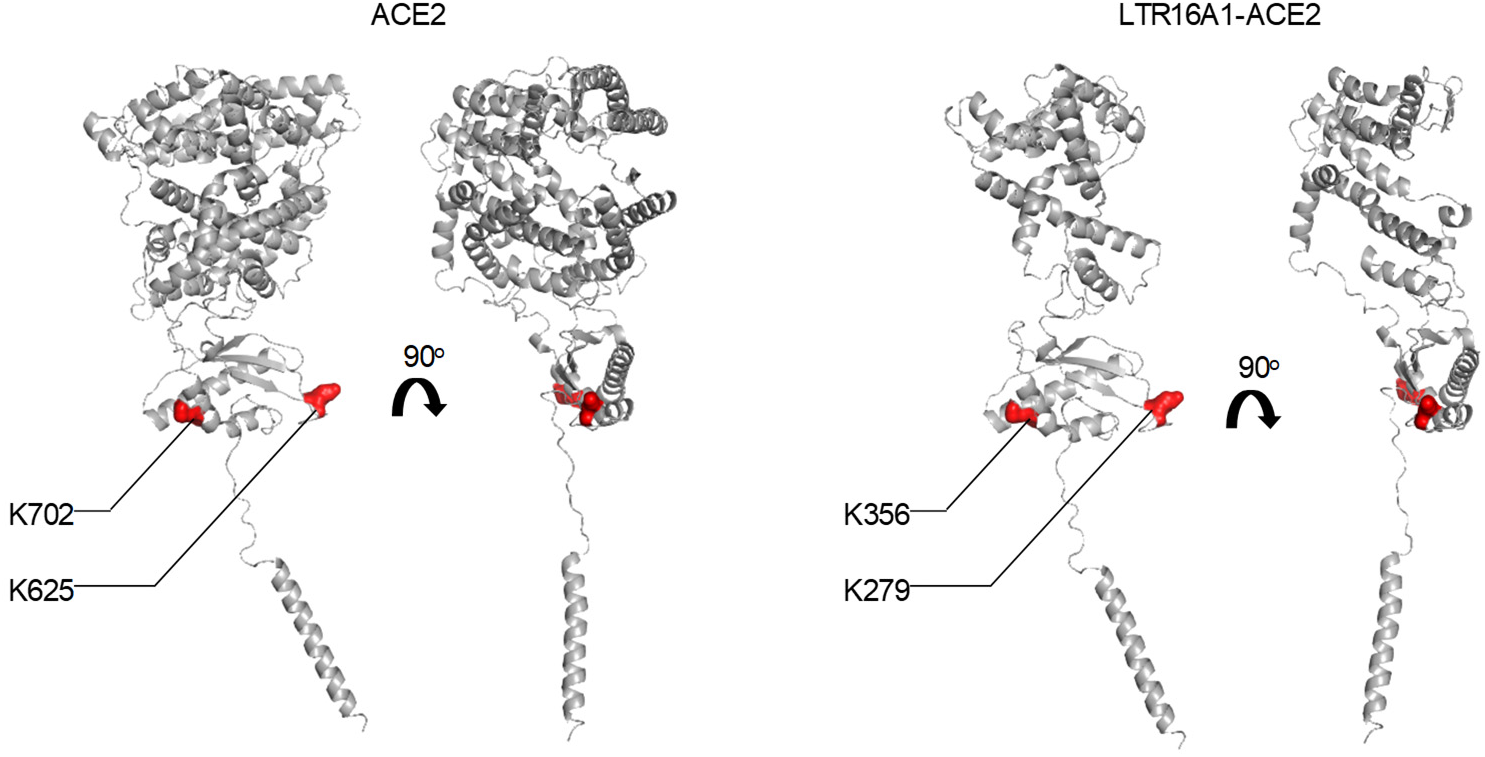
Position of the ubiquitin targets in ACE2 and *LTR16A1-ACE2* protein product. Structure of ACE2 (*left*) and predicted structure of the *LTR16A1-ACE2* protein product (*right*) depicting the position of the two mutated K residues, targeted for ubiquitination.

## References

Attig, J., Young, G.R., Hosie, L., Perkins, D., Encheva-Yokoya, V., Stoye, J.P., Snijders, A.P., Ternette, N., and Kassiotis, G. (2019). LTR retroelement expansion of the human cancer transcriptome and immunopeptidome revealed by de novo transcript assembly. Genome research 29, 1578–1590.

Attig, J., Young, G.R., Stoye, J.P., and Kassiotis, G. (2017). Physiological and Pathological Transcriptional Activation of Endogenous Retroelements Assessed by RNA-Sequencing of B Lymphocytes. Frontiers in microbiology 8, 2489.

Blanco-Melo, D., Nilsson-Payant, B.E., Liu, W.C., Uhl, S., Hoagland, D., Møller, R., Jordan, T.X., Oishi, K., Panis, M., Sachs, D., et al. (2020). Imbalanced Host Response to SARS-CoV-2 Drives Development of COVID-19. Cell 181, 1036–1045.e1039.

Burns, K.H., and Boeke, J.D. (2012). Human transposon tectonics. Cell 149, 740–752.

Chuong, E.B., Elde, N.C., and Feschotte, C. (2016). Regulatory evolution of innate immunity through co-option of endogenous retroviruses. Science (New York, NY) 351, 1083–1087.

Feschotte, C., and Gilbert, C. (2012). Endogenous viruses: insights into viral evolution and impact on host biology. Nat Rev Genet 13, 283–296.

García-Sastre, A. (2017). Ten Strategies of Interferon Evasion by Viruses. Cell host & microbe 22, 176–184.

Gibbert, K., Schlaak, J.F., Yang, D., and Dittmer, U. (2013). IFN-α subtypes: distinct biological activities in anti-viral therapy. British journal of pharmacology 168, 1048–1058.

Hamming, I., Cooper, M.E., Haagmans, B.L., Hooper, N.M., Korstanje, R., Osterhaus, A.D., Timens, W., Turner, A.J., Navis, G., and van Goor, H. (2007). The emerging role of ACE2 in physiology and disease. The Journal of pathology 212, 1–11.

Han, X., Zhou, Z., Fei, L., Sun, H., Wang, R., Chen, Y., Chen, H., Wang, J., Tang, H., Ge, W., et al. (2020). Construction of a human cell landscape at single-cell level. Nature 581, 303–309.

Hoffmann, M., Kleine-Weber, H., Schroeder, S., Kruger, N., Herrler, T., Erichsen, S., Schiergens, T.S., Herrler, G., Wu, N.H., Nitsche, A., et al. (2020). SARS-CoV-2 Cell Entry Depends on ACE2 and TMPRSS2 and Is Blocked by a Clinically Proven Protease Inhibitor. Cell 181, 271–280.e278.

Hung, I.F., Lung, K.C., Tso, E.Y., Liu, R., Chung, T.W., Chu, M.Y., Ng, Y.Y., Lo, J., Chan, J., Tam, A.R., et al. (2020). Triple combination of interferon beta-1b, lopinavir-ritonavir, and ribavirin in the treatment of patients admitted to hospital with COVID-19: an open-label, randomised, phase 2 trial. Lancet (London, England) 395, 1695–1704.

Ivashkiv, L.B., and Donlin, L.T. (2014). Regulation of type I interferon responses. Nature reviews Immunology 14, 36–49.

Kassiotis, G., and Stoye, J.P. (2016). Immune responses to endogenous retroelements: taking the bad with the good. Nat Rev Immunol 16, 207–219.

Kim, D., Paggi, J.M., Park, C., Bennett, C., and Salzberg, S.L. (2019). Graph-based genome alignment and genotyping with HISAT2 and HISAT-genotype. Nature biotechnology 37, 907–915.

Kopecky-Bromberg, S.A., Martínez-Sobrido, L., Frieman, M., Baric, R.A., and Palese, P. (2007). Severe acute respiratory syndrome coronavirus open reading frame (ORF) 3b, ORF 6, and nucleocapsid proteins function as interferon antagonists. Journal of virology 81, 548–557.

Major, J., Crotta, S., Llorian, M., McCabe, T.M., Gad, H.H., Priestnall, S.L., Hartmann, R., and Wack, A. (2020). Type I and III interferons disrupt lung epithelial repair during recovery from viral infection. Science (New York, NY).

Ng, K.W., Attig, J., Young, G.R., Ottina, E., Papamichos, S.I., Kotsianidis, I., and Kassiotis, G. (2019). Soluble PD-L1 generated by endogenous retroelement exaptation is a receptor antagonist. eLife 8.

Patro, R., Duggal, G., Love, M.I., Irizarry, R.A., and Kingsford, C. (2017). Salmon provides fast and bias-aware quantification of transcript expression. Nature methods 14, 417–419.

Sadler, A.J., and Williams, B.R. (2008). Interferon-inducible antiviral effectors. Nature reviews Immunology 8, 559–568.

Shang, J., Ye, G., Shi, K., Wan, Y., Luo, C., Aihara, H., Geng, Q., Auerbach, A., and Li, F. (2020). Structural basis of receptor recognition by SARS-CoV-2. Nature 581, 221–224.

Singh, M., Bansal, V., and Feschotte, C. (2020). A single-cell RNA expression map of human coronavirus entry factors. bioRxiv, 2020.2005.2008.084806.

Stetson, D.B., and Medzhitov, R. (2006). Type I interferons in host defense. Immunity 25, 373–381.

Stukalov, A., Girault, V., Grass, V., Bergant, V., Karayel, O., Urban, C., Haas, D.A., Huang, Y., Oubraham, L., Wang, A., et al. (2020). Multi-level proteomics reveals host-perturbation strategies of SARS-CoV-2 and SARS-CoV. bioRxiv, 2020.2006.2017.156455.

Tange, O. (2011). GNU Parallel: The Command-Line Power Tool. The USENIX Magazine 36, 42–47.

Thompson, P.J., Macfarlan, T.S., and Lorincz, M.C. (2016). Long Terminal Repeats: From Parasitic Elements to Building Blocks of the Transcriptional Regulatory Repertoire. Molecular cell 62, 766–776.

Thorvaldsdóttir, H., Robinson, J.T., and Mesirov, J.P. (2013). Integrative Genomics Viewer (IGV): high-performance genomics data visualization and exploration. Briefings in bioinformatics 14, 178–192.

Tokuyama, M., Kong, Y., Song, E., Jayewickreme, T., Kang, I., and Iwasaki, A. (2018). ERVmap analysis reveals genome-wide transcription of human endogenous retroviruses. Proceedings of the National Academy of Sciences of the United States of America 115, 12565–12572.

Wang, N., Zhan, Y., Zhu, L., Hou, Z., Liu, F., Song, P., Qiu, F., Wang, X., Zou, X., Wan, D., et al. (2020). Retrospective Multicenter Cohort Study Shows Early Interferon Therapy Is Associated with Favorable Clinical Responses in COVID-19 Patients. Cell host & microbe.

Young, G.R., Eksmond, U., Salcedo, R., Alexopoulou, L., Stoye, J.P., and Kassiotis, G. (2012). Resurrection of endogenous retroviruses in antibody-deficient mice. Nature 491, 774–778.

Young, G.R., Mavrommatis, B., and Kassiotis, G. (2014). Microarray analysis reveals global modulation of endogenous retroelement transcription by microbes. Retrovirology 11, 59.

Ziegler, C.G.K., Allon, S.J., Nyquist, S.K., Mbano, I.M., Miao, V.N., Tzouanas, C.N., Cao, Y., Yousif, A.S., Bals, J., Hauser, B.M., et al. (2020). SARS-CoV-2 Receptor ACE2 Is an Interferon-Stimulated Gene in Human Airway Epithelial Cells and Is Detected in Specific Cell Subsets across Tissues. Cell 181, 1016–1035.e1019.

